# ChineseEEG: A Chinese Linguistic Corpora EEG Dataset for Semantic Alignment and Neural Decoding

**DOI:** 10.1101/2024.02.08.579481

**Authors:** Xinyu Mou, Cuilin He, Liwei Tan, Junjie Yu, Huadong Liang, Jianyu Zhang, Tian Yan, Yu-Fang Yang, Ting Xu, Qing Wang, Miao Cao, Zijiao Chen, Chuan-Peng Hu, Xindi Wang, Quanying Liu, Haiyan Wu

## Abstract

An Electroencephalography (EEG) dataset utilizing rich text stimuli can advance the understanding of how the brain encodes semantic information and contribute to semantic decoding in brain-computer interface (BCI). Addressing the scarcity of EEG datasets featuring Chinese linguistic stimuli, we present the ChineseEEG dataset, a high-density EEG dataset complemented by simultaneous eye-tracking recordings. This dataset was compiled while 10 participants silently read approximately 11 hours of Chinese text from two well-known novels. This dataset provides long-duration EEG recordings, along with pre-processed EEG sensor-level data and semantic embeddings of reading materials extracted by a pre-trained natural language processing (NLP) model. As a pilot EEG dataset derived from natural Chinese linguistic stimuli, ChineseEEG can significantly support research across neuroscience, NLP, and linguistics. It establishes a benchmark dataset for Chinese semantic decoding, aids in the development of BCIs, and facilitates the exploration of alignment between large language models and human cognitive processes. It can also aid research into the brain’s mechanisms of language processing within the context of the Chinese natural language.

## Background & Summary

The human brain’s ability to rapidly comprehend linguistic information and generate corresponding linguistic expressions is an indicator of its complex processing capabilities^1^. When exposed to linguistic stimuli, the human brain encodes the semantic information through neural activities^2^. By analyzing such neural activities, we can uncover the encoding mechanisms of semantics in the brain^3^. A variety of neural signals, including EEG, Functional Magnetic Resonance Imaging (fMRI), Electrocorticography (ECoG) are employed in language-related tasks, from academic research like investigating language processing in the brain to practical applications like language decoding in BCI^4–9^. Recently, a lot of studies on neurolinguistics utilized both machine learning methods and modern deep learning methods in NLP to explore linguistic-related problems^10–16^. However, these data-driven methods rely heavily on massive and comprehensive datasets^17^. In the field of NLP, it is relatively easy to collect large amounts of natural language data. In contrast, acquiring a large volume of neural signals generated in response to natural language stimuli poses significant challenges. To utilize the strong ability of modern data-driven methods, it is important to scale neural datasets to commensurate the state-of-the-art NLP to encompass the wide range of language expressions encountered in daily life. Among all neuroimaging techniques, EEG holds great potential to meet this demand. EEG is non-invasive and cost-effective^18^, which allows the creation of long-duration neural signal datasets enriched with semantic information. Meanwhile, EEG features high temporal resolution^19^, which enables it to precisely capture the brain’s rapid dynamic changes in the language processing process.

Despite the abundance of EEG datasets for natural visual stimuli (e.g., THINGS-EEG)^20–23^, those for natural language stimuli remain scarce. Currently, only a few language-related EEG datasets exist, such as the ZuCo dataset^24^. However, the majority of these datasets are collected using stimuli from English language corpora. This leads to limited research on the neural representations of other languages like Chinese. The brain’s processing mechanisms differ for various languages. For example, the brain exhibits specificity in response to Chinese compared to English^25^. Therefore, it is important to create an EEG dataset based on other language stimuli. Chinese, being distinct from English in both structure and semantics, provides an opportunity to expand our understanding of neural responses to linguistic stimuli. An EEG dataset stimulated by Chinese corpora can facilitate the investigation of cross-linguistic commonalities and variations in language processing in the brain, bringing new perspectives to our understanding of language processing mechanisms.

To address these gaps, we have collected an EEG dataset, named the “ChineseEEG” (Chinese Linguistic Corpora EEG Dataset). It contains high-density EEG data and simultaneous eye-tracking data recorded from 10 participants, each silently reading Chinese text for about 11 hours. The text materials are sourced from two well-known novels, *The Little Prince* and *Garnett Dream*, both in their Chinese versions. This dataset further comprises multiple versions of pre-processed EEG sensor-level data generated under different parameter settings, offering researchers a diverse range of selections. Additionally, we provide embeddings of the Chinese text materials encoded from BERT-base-chinese model, which is a pre-trained NLP model specifically used for Chinese ^26^, aiding researchers in exploring the alignment between text embeddings from NLP models and brain information representations in neural signals.

ChineseEEG is a pilot EEG dataset specifically stimulated by Chinese text. It offers several advantages. Firstly, each participant was exposed to around 11 hours of diverse Chinese linguistic stimuli, encompassing a broad spectrum of semantic information. The extensive exposure is significant for studying the long-term neural dynamics of language processing in the brain. Secondly, we employed 128 channels of high-density EEG data, which offers superior spatial resolution for precise localization of brain regions involved in language processing. Besides, with a sampling rate of 1 kHz, it effectively captures the dynamics of neural representations during reading. Thirdly, EEG data generated from Chinese language stimuli will significantly support research within the Chinese context, aiding researchers in revealing the characteristics of brain signal representations under Chinese stimuli, and promoting the development of brain-to-text translation, semantic decoding and other practical applications tailored to Chinese context. This dataset can also bring diversity to languages used in related research, encouraging the exploration of similarities and differences in language processing stimulated by different languages. Lastly, this dataset can effectively facilitate the integration of neuroscience and computer science methodologies. The inclusion of the text embeddings is beneficial for scholars in neuroscience domain who lack text processing experience, enabling them to directly utilize the embeddings from computational linguistic models to explore neuroscience problems. The dataset can also facilitate the entry of computer science scholars into the field of neuroscience, enabling them to use computational methods to explore topics in neuroscience such as the encoding mechanisms of the Chinese language in the brain and the utilization of EEG for text decoding.

## Methods

### Participants and task overview

We recruited 15 participants (18-26 years old, averaged 21.26 years old, and 8 males). 3 participants participated the pre-experimental test before the official experiment to ensure the rationality of the experimental procedure and the stability of the devices. In the official experiment, 2 participants withdrew halfway due to scheduling conflicts. In total, data from only 10 participants were used (18-24 years old, averaged 20.68 years old, and 5 males). No participant reported neurological or psychiatric history. All participants are right-handed and have normal or corrected-to-normal vision. Each participant voluntarily enrolled in and signed the informed consent form before the experiment and got a coupon compensation of approximately 50 MOP (MOP is the official currency of the Macao Special Administrative Region of China) for each experimental run (25 runs in total). This study complied with the Declaration of Helsinki and was performed according to the ethics committee approval of the Institutional Review Board of the University of Macau.

### Experimental material

The experimental materials consist of two novels, both in the genre of children’s literature. The first is the Chinese translation of *The Little Prince* and the second is *Garnett Dream*. The text of these novels was sourced from the internet. Using novels, especially children’s literature provides several advantages for research, especially within a naturalistic paradigm. Firstly, given their extensive size, these novels offer vast and diverse linguistic content, encompassing the majority of frequently utilized Chinese characters and daily expressions. Besides, children’s literature can create an engaging environment for participants, making them more focused and emotionally engaged in the experiment.

Each novel was used as the material for a single session in the experiment. Each session was divided into several runs. For *The Little Prince*, the preface was used as the material for the practice reading phase. The main body of the novel was then used for seven runs in the formal reading phase. The first six runs each includes 4 chapters of the novel, while the seventh run includes the last two chapters. For *Garnett Dream*, the first 18 chapters were used for 18 runs in the formal reading stage, with each run including a complete chapter. Due to the loss of markers during the EEG collection process, run 18 of ses-GarnettDream of sub-07 is unusable. We request this participant to re-complete the reading task using Chapter 19 of *Garnett Dream*.

To properly present the text on the screen during the experiment, the content of each run was segmented into a series of units, with each unit containing no more than 10 Chinese characters. These segmented contents were saved in Excel (.xlsx) format for subsequent usage. During the experiment, three adjacent units from each run’s content will be displayed on the screen in three separate lines, with the middle line highlighted for the participant to read. The relevant code has been uploaded to the GitHub repository. See Code availability section for detailed information.

The overview of experimental materials is shown in Table 1. In summary, a total of 115,233 characters (24,324 in *The Little Prince* and 90,909 in *Garnett Dream*), of which 2,985 characters are unique, are used as experimental stimuli in ChineseEEG dataset.

**Table 1.** An overview of the experiment.

| Session | Run | Chapter | Number of Chinese characters | Duration |
| --- | --- | --- | --- | --- |
| LittlePrince |  | Preface | 210 |  |
|  | 1 | 1-4 | 3,805 | 24min34s |
|  | 2 | 5-8 | 3,734 | 24min5s |
|  | 3 | 9-12 | 3,218 | 20min50s |
|  | 4 | 13-16 | 4,030 | 25min59s |
|  | 5 | 17-20 | 1,713 | 11min11s |
|  | 6 | 21-24 | 3,635 | 23min27s |
|  | 7 | 25-27 | 4,189 | 26min54s |
| GarnettDream | 1 | 1 | 5,267 | 34min17s |
|  | 2 | 2 | 4,406 | 28min39s |
|  | 3 | 3 | 5,327 | 34min35s |
|  | 4 | 4 | 3,906 | 25min15s |
|  | 5 | 5 | 4,989 | 32min14s |
|  | 6 | 6 | 4,413 | 28min29s |
|  | 7 | 7 | 3,912 | 25min25s |
|  | 8 | 8 | 5,537 | 35min52s |
|  | 9 | 9 | 4,171 | 27min2s |
|  | 10 | 10 | 5,943 | 38min30s |
|  | 11 | 11 | 4,351 | 28min21s |
|  | 12 | 12 | 4,830 | 31min13s |
|  | 13 | 13 | 3,799 | 24min31s |
|  | 14 | 14 | 4,963 | 32min9s |
|  | 15 | 15 | 4,656 | 29min55s |
|  | 16 | 16 | 4,615 | 29min42s |
|  | 17 | 17 | 5,273 | 33min57s |
|  | 18 | 18 | 5,113 | 32min57s |
|  | 19 | 19 | 5,438 | 35min10s |

### Experimental procedures

Participants were instructed to sit in an adjustable chair, whose eyes were approximately 67 cm away from the monitor (Dell, width: 54 cm, height: 30.375 cm, resolution: 1,920×1,080 pixels, vertical refresh rate: 60 Hz), see Figure 1b. They were tasked with reading a novel and were required to keep their heads still and keep their gaze on the highlighted (red) Chinese characters moving across the screen, reading at a pace set by the program. They were required to read an entire novel in multiple runs within a single session. Each run is divided into two phases: the eye-tracker calibration phase and the reading phase, with a break between two adjacent runs to allow the experimenter to check the electrodes’ impedance and add saline if necessary. Each run includes either 3 to 4 chapters of *The Little Prince* or a single chapter of *Garnett Dream*, lasting approximately 30 minutes.

**Figure 1.**
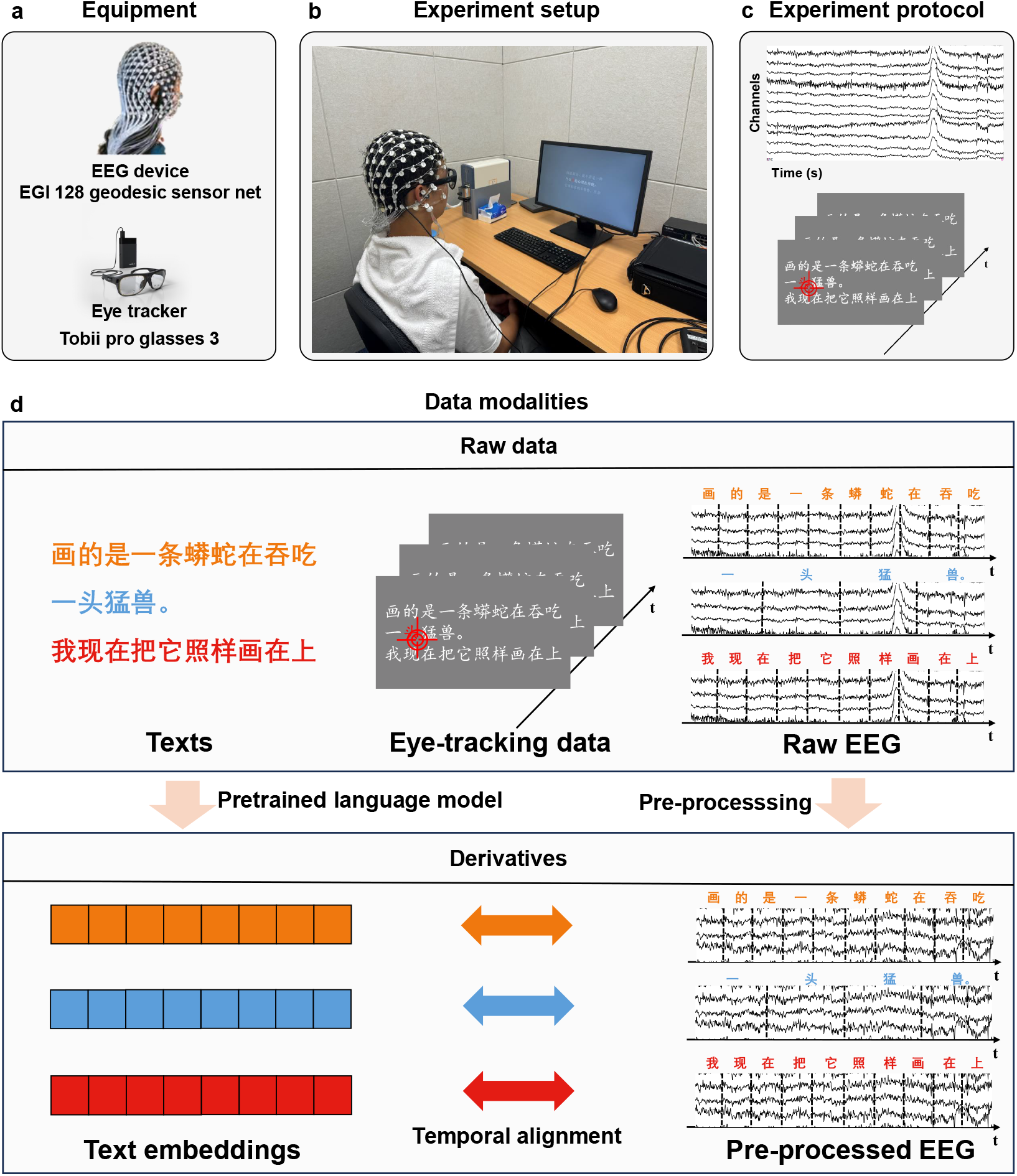
Overview of the experiment and the modalities included in the dataset. (a) Equipment utilized in the experiment, including the EGI device for collecting EEG data and the Tobii Glasses 3 eye-tracker for tracking eye movements. (b) The experiment setup. Participants were instructed to sit quietly approximately 67cm from the screen and sequentially read the highlighted text. (c) The experimental protocol. Participants’ 128-channel EEG signals and eye-tracking data were recorded while reading the highlighted text. (d) The data modalities in the dataset. The dataset comprises raw data such as the original textual stimuli, eye movement data, EEG data, and derivatives such as text embeddings from pre-trained NLP models and pre-processed EEG data.

The presentation of stimuli was managed using PsychoPy v2023.2.3^27^, with the EGI PyNetstation v1.0.1 module facilitating the connection between PsychoPy and EGI Netstation. We also utilized g3pylib package to control our eye-tracker to follow the eye movement trajectories of the participants.

#### Phase 1: Eye-tracker calibration phase

At the beginning of each run, participants were required to undergo an eye-tracker calibration process. Initially, the message “Hello! Please press the spacebar to start calibration” was displayed at the screen’s center. Participants were instructed to keep their gaze at a fixation point, which sequentially appeared at the four corners and the center of the screen, each for 5 seconds. If the calibration failed, participants were prompted to start another calibration. Upon successful calibration, the message “Calibration successful! The page will automatically redirect in 5 seconds” was displayed at the center of the screen.

#### Phase 2: Reading phase

After the calibration phase, participants were automatically directed to the reading phase. During the reading process, the screen initially displayed the serial number of the current chapter. Subsequently, the text appeared with three lines per page, ensuring each line contained no more than ten Chinese characters (excluding punctuation). On each page, the middle line was highlighted as the focal point, while the upper and lower lines were displayed with reduced intensity as the background. Each character in the middle line was sequentially highlighted with red color for 0.35 s, and participants were required to read the novel content following the highlighted cues.

It should be noted that during the initial participation in the experiment, participants were required to complete a practice reading phase. The preface chapter of *The Little Prince* was selected as the reading material for this phase. All settings remained the same as those of the formal reading stage, to familiarize participants with the eye-tracker calibration process and the reading task.

After each run, participants were provided with adequate rest time until they reported ready to start the subsequent run. During the rest period, the experimenter replenished the saline solution on the electrodes of the EEG cap, which helped to maintain a low impedance, ensuring the collection of high-quality EEG data. Additionally, the experimenter checked the power status of the eye-tracker and replaced the batteries as necessary to ensure its continuous operation.

### Data collection and analysis

This section shows the details of the data collection, pre-processing, and data analysis procedure. The modalities included in our dataset are shown in 1d, including raw data and derivatives. Raw data contains the raw EEG data, eye-tracking data, raw text materials, and derivatives contain pre-processed EEG data and text embeddings generated by a pre-trained NLP model BERT-base-chinese.

#### EEG data collection

EEG data was acquired using an EGI 128-channel cap based on the GSN-HydroCel-128 montage with the Geodesic Sensor Net system (see Figure 1a). The egi-pynetstation v1.0.1 package was used to control the EGI system. During recording, the sampling rate was 1 kHz. The impedance of each electrode was kept below 50 kΩ during the experiment. Setups and recording parameters are similar to our previous EEG dataset^28^. To precisely co-register EEG segments with individual characters during the experiment, we marked the EEG data with triggers (Table 2). The raw EEG data was exported to metafile format (.mff) files on the macOS system.

**Table 2.** EEG triggers.

| Trigger | Description |
| --- | --- |
| EYES | Start of eye-tracker recording |
| EYEE | End of eye-tracker recording |
| CALS | Start of the calibration stage before reading |
| CALE | End of the calibration stage |
| BEGN | Start of EEG data collection by the EGI device |
| STOP | Stop collecting EEG data |
| CHxx | Start of each chapter, where xx is the chapter number (e.g., the first chapter is CH01) |
| ROWS | Start of a new line of text |
| ROWE | End of a line |
| PRES | Start of the preface reading phase |
| PREE | End of the preface reading phase |

#### Eye-tracking data collection

Eye-tracking data was acquired using Tobii Pro Glasses 3 (see Figure 1a). The device features 16 illuminators and 4 eye cameras integrated into scratch-resistant lenses, along with a wide-angle scene camera, allowing for a comprehensive capture of participant behavior and environmental context. We utilized the package g3pylib to control the glasses. During recording, the sampling rate was set to 100 Hz. The raw data was exported to .zip files.

#### EEG data pre-processing

To retain maximum amount of valid information in the data, we performed minimal pre-processing on the data, allowing researchers to further process the data according to their specific research needs. The pre-processing pipeline is shown in Figure 2. These pre-processing steps include data segmentation, downsampling, powerline filtering, band-pass filtering, bad channel interpolation, independent component analysis (ICA), and re-referencing. The MNE v1.6.0^29^ package was utilized to implement all pre-processing steps.

**Figure 2.**
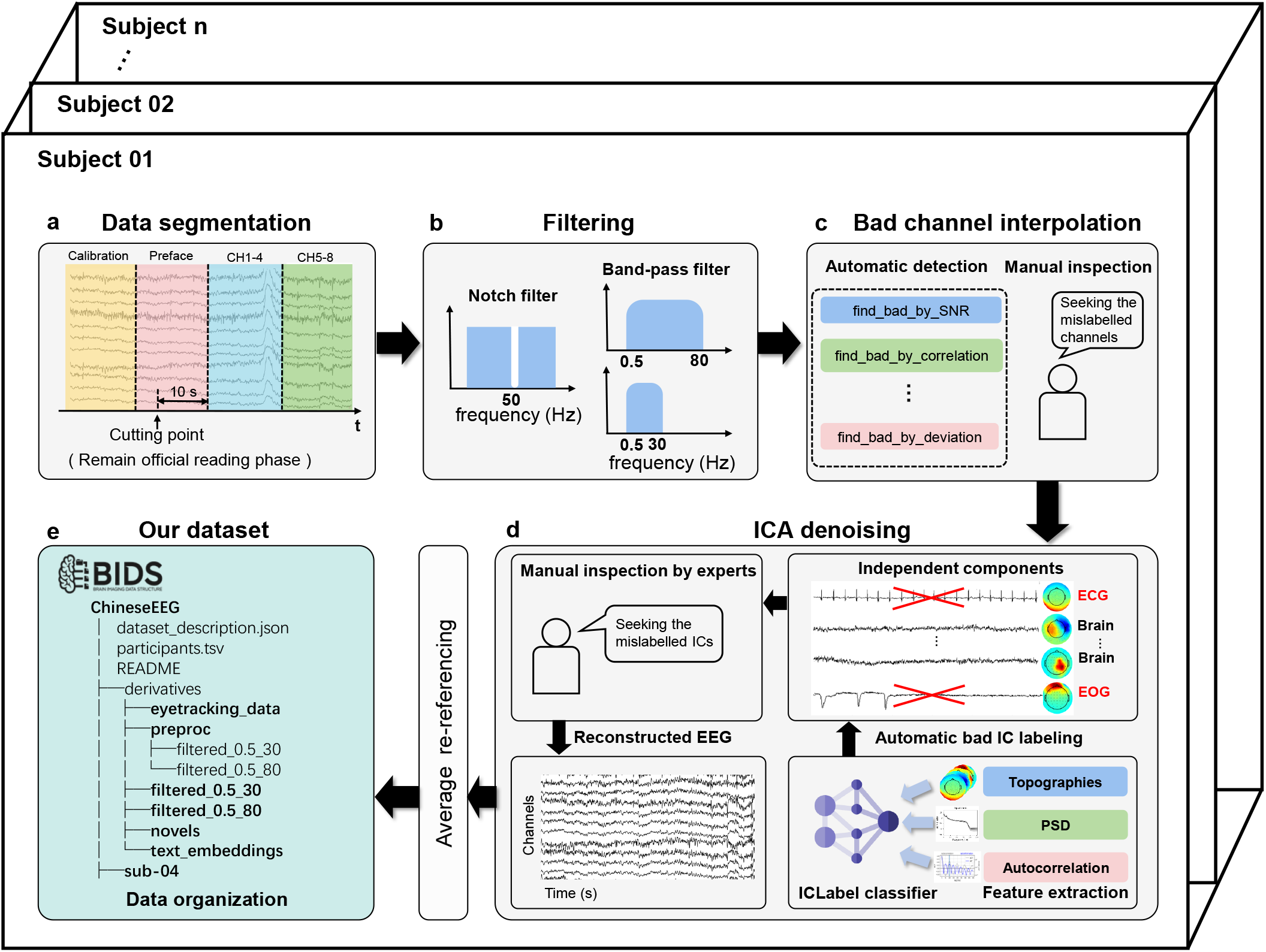
EEG pre-processing pipeline. (a) Data segmentation: Data is segmented based on markers, retaining only the data from the formal reading phase. (b) Band-pass filtering: Two versions of filtered data are provided, with band-pass ranges of 0.5-30 Hz and 0.5-80 Hz respectively. (c) Bad channel interpolation: Our bad channel detection includes automatic detection implemented with the pyprep package and manual checking. For interpolation, the spherical spline interpolation implemented in MNE is utilized. (d) ICA denoising: In this part, the automatic labeling method in mne-iclabel package is utilized followed by a manual checking to remove noisy independent components such as eye movements and heartbeats. (e) Dataset organization: Our dataset is organized in the BIDS format. The detailed file structure is shown in Figure 3.

During the data segmentation phase, we only retained data from the formal reading phase of the experiment. Based on the event markers during the data collection phase, we segmented the data, removing sections irrelevant to the formal experiment such as calibration and preface reading. To minimize the impact of subsequent filtering steps on the beginning and end of the signal, an additional 10 seconds of data was retained before the start of the formal reading phase. Subsequently, the signal was downsampled to 256 Hz.

Following this, a 50 Hz notch filter was applied to remove the powerline noise from the signal. Next, we performed band-pass overlap-add FIR filter on the signal to eliminate the low-frequency direct current components and high-frequency noise. Here, two versions of filtered data were offered. The first one has a filter band of 0.5-80 Hz and the second one has a filter band of 0.5-30 Hz. Researchers can choose the appropriate version based on their specific needs. After filtering, we performed an interpolation of bad channels. The bad channels were selected automatically using a Python-implemented EEG pre-processing package pyprep v0.4.3

. After automatic detection, we manually checked to avoid mislabeling or errors before interpolation. The spherical spline interpolation in the MNE package was utilized in this process.

Independent Component Analysis (ICA) was then applied to the data, utilizing the infomax algorithm available in the MNE package. The number of independent components was set to 20, ensuring that they contain the majority of information while not being so numerous to increase the burden of manual processing. Additionally, we set the random seed of the ICA algorithm to 97 to ensure the reproducibility of the ICA results. An automatic method was used to inspect and label components. It was implemented using mne-iclabel v0.5.1^30^, which is a Python-implemented package for automatic independent component labeling. By manually inspecting the independent components after automatic labeling, we excluded obvious noise components such as Electrooculography (EOG) and Electrocardiogram (ECG). Finally, the data was re-referenced using the average method.

The process of manually identifying bad channels and excluding independent components during the ICA step can be conducted through annotations in a Graphical User Interface (GUI), making the annotation process quicker and more user-friendly.

## Data Records

The full dataset is publicly accessible via the ChineseNeuro Symphony community (CHNNeuro) in the Science Data Bank (ScienceDB) platform (https://doi.org/10.57760/sciencedb.CHNNeuro.00002) or via the Openneuro platform (https://openneuro.org/datasets/ds004952).

### EEG data organization

The dataset is organized following the EEG-BIDS^31^ specification, which is an extension to the brain imaging data structure for EEG. The overview directory tree of our dataset is shown in Figure 3. The dataset contains some regular BIDS files, 10 participants’ data folders, and a *derivatives* folder. The stand-alone files offer an overview of the dataset: i) *dataset_description.json* is a JSON file depicting the information of the dataset, such as the name, dataset type and authors; ii) *participants.tsv* contains participants’ information, such as age, sex, and handedness; iii) *participants.json* describes the column attributes in *participants.tsv*; iv) *README.md* contains a detailed introduction of the dataset.

**Figure 3.**
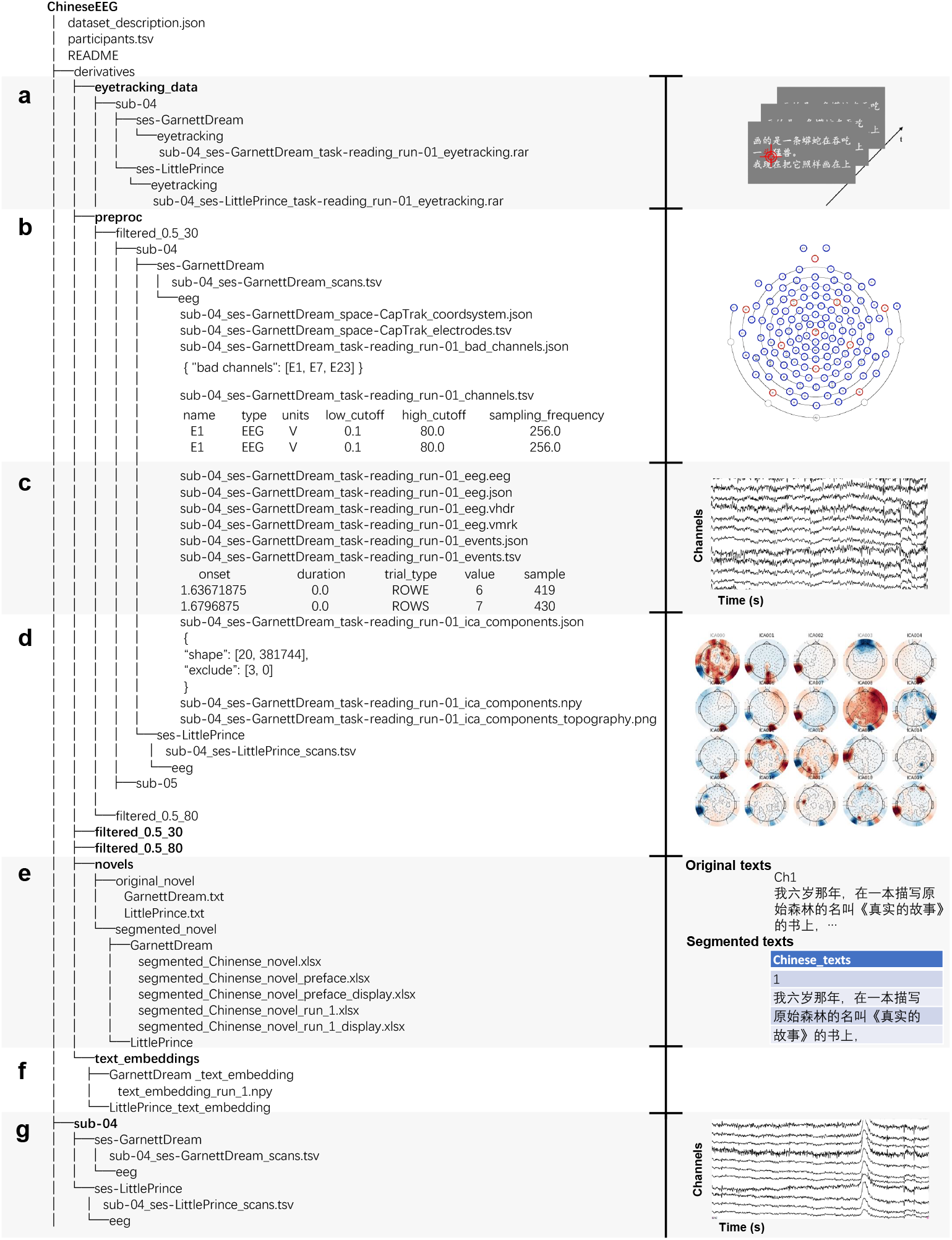
File structure of the dataset. (a) Eye-tracking data: Each experimental run is associated with a .zip file that contains eye-tracking data. (b) Electrode information files: These include detailed information of electrodes such as the location, type, and sampling rate, as well as information on any channels marked as bad during pre-processing. (c) EEG data and event-related files: Including EEG data in BrainVision format and event files that record marker information. (d) ICA-related files: Containing independent components in numpy format, records of removed components during pre-processing, and topographic maps of the components. (e) Text materials: Containing original and segmented text. (f) Text embedding files: Each file corresponds to an experimental run and is stored in .npy format. (g) Raw EEG data.

Each participant’s folder contains two folders named *ses-LittlePrince* and *ses-GarnettDream*, which store the data of this participant reading two novels, respectively. Each of the two folders contains a folder *eeg* and one file *sub-xx_scans.tsv*. The tsv file contains information about the scanning time of each file. The *eeg* folder contains the source raw EEG data of several runs, channels, and marker events files. Each run includes an *eeg.json* file, which encompasses detailed information for that run, such as the sampling rate and the number of channels. Events are stored in *events.tsv* with onset and event ID. The EEG data is converted from raw metafile format (.*mff* file) to BrainVision format (.*vhdr*, .*vmrk* and .*eeg* files) since EEG-BIDS is not officially compatible with .*mff* format. All data is formatted to EEG-BIDS using the mne-bids v0.14^31,32^ package in Python.

The *derivatives* folder contains six folders: *eyetracking_data, filtered_0.5_80, filtered_0.5_30, preproc, novels*, and *text_embeddings*. The *eyetracking_data* folder contains all the eye-tracking data. Each eye-tracking data is formatted in a .*zip* file with eye moving trajectories and other parameters like sampling rate saved in different files. The *filtered_0.5_80* folder and *filtered_0.5_30* folder contain data that has been processed up to the pre-processing step of 0.5-80 Hz and 0.5-30 Hz band-pass filtering respectively. This data is suitable for researchers who have specific requirements and want to perform customized processing on subsequent pre-processing steps like ICA and re-referencing. The *preproc* folder contains minimally pre-processed EEG data that is processed using the whole pre-processing pipeline. It includes four additional types of files compared to the participants’ raw data folders in the root directory: i) *bad_channels.json* contains bad channels marked during bad channel rejection phase. ii) *ica_components.npy* stores the values of all independent components in the ICA phase. iii) *ica_components.json* includes the independent components excluded in ICA (the ICA random seed is fixed, allowing for reproducible results). iv) *ica_components_topography.png* is a picture of the topographic maps of all independent components, where the excluded components are labeled in grey. The *novels* folder contains the original and segmented text stimuli materials. The original novels are saved in .txt format and the segmented novels corresponding to each experimental run are saved in Excel (.xlsx) files. The *text_embeddings* folder contains embeddings of the two novels. The embeddings corresponding to each experimental run are stored in NumPy (.npy) files.

## Technical Validation

### Classic sensor-level EEG analysis

The EEG data in the dataset can be used to do classic time-frequency analysis. In this section, pre-processed EEG data was used to extract neural oscillations in different frequency bands. Specifically, we targeted the segment corresponding to the sentence “Draw me a sheep” in *The Little Prince* from the 0.5-80 Hz filtered pre-processed data of sub-07. The analysis was exclusively focused on the C3 electrode to investigate the neural activities at the scalp location overlying the temporal lobe, which is a language processing related area.

To dissect the frequency components inherent in the C3 electrode’s signal, we applied the Fast Fourier Transform (FFT) algorithm to the data. This mathematical technique transforms the time-domain signal into the frequency domain, revealing the spectrum of frequencies present in the neural recordings. We defined frequency bands of interest—Theta (4-8 Hz), Alpha (8-12 Hz), Beta (12-30 Hz), and Gamma (30-100 Hz)—to categorize the neural oscillations according to their respective frequency ranges.

For each frequency band, we separated the components from the FFT results and conducted an inverse FFT to retrieve the time-domain signal representing the band’s oscillatory activity. This step allows for the quantitative analysis of the amplitude of oscillations within each frequency band, offering insights into the neurophysiological activity in these specific ranges. The results of different frequency bands are shown in Figure 4.

**Figure 4.**
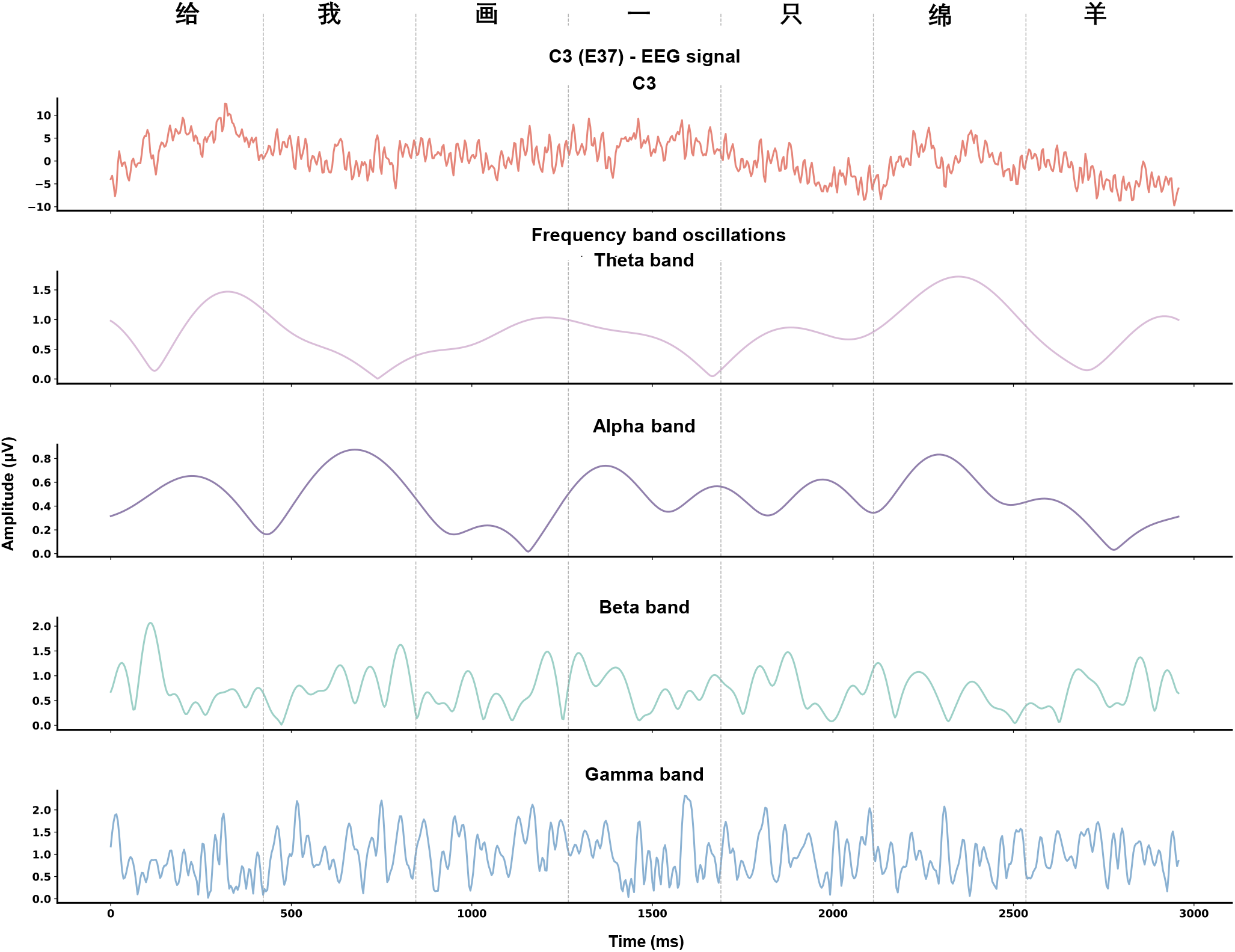
EEG time course and the neural oscillations under different frequency bands (i.e., Theta, Alpha, Beta, and Gamma) corresponding to the Chinese sentence meaning “Draw me a sheep”. The pre-processed EEG data using 0.5-80 Hz band-pass filter from ses-LittlePrince of sub-07 was used in the analysis. We illustrated the EEG signals from electrode C3, which locates at a language processing related area overlying the temporal lobe.

### EEG source reconstruction

Apart from the sensor level analysis, the EEG data allows for conducting source localization. Here, a segment of the data was utilized as an example to perform the source-level analysis using the MNE package. In surface reconstruction, we utilized the fsaverage MRI template in MNE package. A 3-layer Boundary Element Method (BEM) model with 15360 triangles and conductivities of 0.3 S/m, 0.006 S/m, and 0.3 S/m for the brain, skull, and scalp compartments respectively was created. Source spaces consisted of 10242 sources per hemisphere. A segment of the pre-processed EEG data with a band-pass frequency band of 0.5-80 Hz corresponding to one line displayed in the experiment was used to calculate the inverse solution. Inverse solutions were calculated using dynamic Statistical Parametric Maps (dSPM). The method was selected because it is widely used by researchers and is representative of currently used methods^33^. We offer the code of source reconstruction in our GitHub repository. See Code availability section for detailed information.

The visualization of the source activities is shown in Figure 5b. Results for the left and right hemispheres are presented separately. The moments of peak activation in the left and right brain regions are chosen for visualization. The source localization results for the first segment reveal a dispersed activation area, encompassing the anterior temporal lobe and temporo-parietal region, which are associated with language comprehension and primary processing^34^. The results of the second segment exhibit more focused activation, particularly near the left middle temporal gyrus, an area (encompassing Wernicke’s area) intimately related to language comprehension^35^. The activation areas for the third segment are localized in the left temporal and frontal lobes, potentially representing high-level stages of language processing, including sentence construction, semantic processing, and language expression^36^. Figure 5c presents plots of source activities over time, derived from 12 sources in the corresponding region with strongest activities. The first two curves in each plot correspond to sources in the left and right hemispheres that reach maximum peak values.

**Figure 5.**
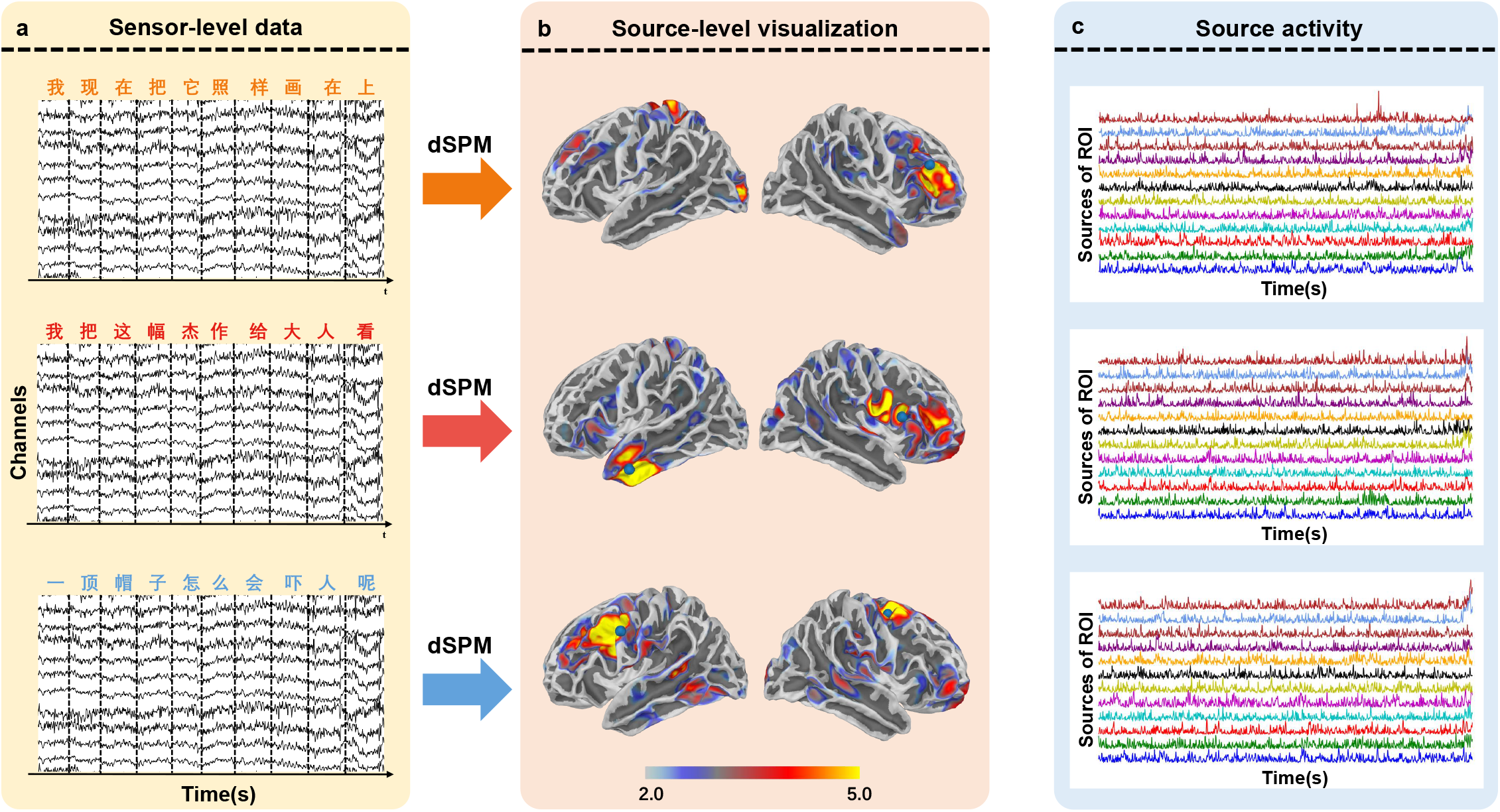
EEG source localization analysis. (a) EEG sensor-level data: Three segments of pre-processed EEG data using 0.5-80 Hz band-pass filter were selected for analysis, accompanied by the corresponding text segments shown above the EEG segments. (b) Visualization of brain activation after source analysis: The dSPM method was utilized to solve the inverse problem. Results for the left and right hemispheres are presented separately. The moments of peak activation in the left and right brain regions are chosen for visualization. (c) Plots of source activity over time: Each plot contains the activities of 12 sources in the region with the strongest activity.

### Text embeddings with pre-trained language model

To assist researchers in efficiently exploring the alignment between EEG and text representations, as well as in text decoding based on EEG, this study provides embeddings of two novels calculated using a pre-trained language model, accompanied by the code to compute these embeddings. This work employed Google’s pre-trained language model BERT-base-Chinese^26^. This model, pre-trained on Chinese corpora, effectively encodes Chinese semantic features. During the experimental procedure, each displayed line of text contains *n* Chinese characters. The BERT-base-Chinese model processes these n Chinese characters, yielding an embedding of size *(n, 768)*, where *n* represents the number of Chinese characters, and *768* the dimensionality of the embedding. To ensure displayed lines of varying length to have embeddings of the same shape, the first dimension of the embeddings is averaged to standardize the embedding size to *(1, 768)* for each instance. This processing procedure was implemented using the Hugging Face Transformers v4.36.2^37^ package.

### Temporal alignment between EEG and text sequences

To facilitate semantic decoding, it is necessary to align specific text with its corresponding EEG segment in the temporal domain. During the marking process when collecting the data, the start and end of each line of the stimuli were annotated, thereby enabling the alignment of each text line with a corresponding segment of EEG data. Given the consistent highlighting duration for each character, the EEG segment can be equally divided to match the corresponding character. In the GitHub repository, we offer the script to align the EEG segments to their corresponding text and text embeddings.

## Usage Notes

### Prior to using the data

The code for the experiment and data analysis has been uploaded to GitHub to facilitate sharing and utilization, which is accessible at https://github.com/ncclabsustech/Chinese_reading_task_eeg_processing.

The code repository contains four main modules, each including scripts desired to reproduce the experiment and data analysis procedures. The script *cut_chinese_novel.py* in the *novel_segmentation_and_text_embeddings* folder contains the code to prepare the stimulation materials from source materials. The script *play_novel.py* in the *experiment* module contains code for the experiment, including text stimuli presentation and control of the EGI device and Tobii Glasses 3 eye-tracker. The script *preprocessing.py* in *data_preprocessing_and_alignment* module contains the main part of the code to apply pre-processing on EEG data. The script *align_eeg_with_sentence.py* in the same module contains code to align the EEG segments with corresponding text contents and text embeddings. The *docker* module contains the Docker image required for deploying and running the code, as well as tutorials on how to use Docker for environment deployment.

The code for EEG data pre-processing is highly configurable, permitting flexible adjustments of various pre-processing parameters, such as data segmentation range, downsampling rate, filtering range, and choice of ICA algorithm, thereby ensuring convenience and efficiency. Researchers can modify and optimize this code according to their specific requirements.

Before using our ChineseEEG dataset, we encourage all users to check the *README.md* and the updated information in the GitHub repository.

### Potentials opportunities

The ChineseEEG dataset is a potential resource for accelerating the exploration of scientific problems such as brain’s neural representations of semantic information, and mechanisms of the human brain in learning, memory, and attention. It can also contribute in enhancing the development of applications such as BCI systems.

The utilization of ChineseEEG dataset can deepen our understanding of the learning process of languages in the human brain, especially how the human brain learns Chinese, such as holistic Chinese word recognition^38^. Besides, This dataset can also help us in exploring representations in EEG that reflect the language processing process, along with their association with brain functions such as decision making, memory storage and retrieval.

The ChineseEEG dataset also offers crucial opportunities in practical applications like brain-to-text BCI. The abundant data in the dataset can facilitate the utilization of modern data-driven methods from NLP in language related tasks, such as training large-scale models to learn the complex semantic patterns in neural signals, and aligning neural signals with natural languages in the representation space. For example, by using large-scale neural data to train deep learning models, these models can effectively learn the complex semantic representations of the brain under linguistic stimuli and generalize well across a wide range of downstream tasks, such as semantic decoding^39^, text-based emotion recognition^40^ and sentiment classification^41^. It can also mitigate the challenge of inter-subject generalization in BCI systems caused by the variability of neural signals among individuals. By training the model on vast neural signals enriched with diverse semantic information from different subjects, the model can learn to extract invariant semantic patterns and structures across individuals, thereby becoming more adaptable to a wide range of individuals.

Given that most existing EEG datasets primarily focus on English language materials, the ChineseEEG dataset can be especially useful for exploring both scientific problems and practical applications in the context of Chinese language, prompting cross-cultural research in related fields.

## Code availability

The code for all modules is openly available on GitHub (https://github.com/ncclabsustech/Chinese_reading_task_eeg_processing). All scripts were developed in Python 3.10^42^. Package openpyxl v3.1.2 was utilized to export segmented text in Excel (.xlsx) files, and egi-pynetstation v1.0.1, g3pylib v0.1.1, psychopy v2023.2.3^27^ were used to implement the scripts for EGI device control, Tobii eye-tracker control, stimuli presentation respectively. In the data pre-processing scripts, MNE v1.6.0^29^, pybv v0.7.5^43^, pyprep v0.4.3^44^, mne-iclabel v0.5.1^30^ were used to implement the pre-processing pipeline, while mne-bids v0.14^31,32^ was used to organize the data into BIDS format. The text embeddings were calculated using Hugging Face transformers v4.36.2^37^. For more details about code usage, please refer to the GitHub repository.

## Acknowledgements

This work was mainly supported by the MindD project of Tianqiao and Chrissy Chen Institute(TCCI), the Science and Technology Development Fund (FDCT) of Macau [0127/2020/A3, 0041/2022/A], the Natural Science Foundation of Guangdong Province(2021A1515012509), Shenzhen-Hong Kong-Macao Science and Technology Innovation Project (Category C) (SGDX2020110309280100), and the SRG of University of Macau (SRG2020-00027-ICI). We also thank all research assistants who provided general support in participant recruiting and data collection.

## Author contributions statement

H.Wu, Q.Liu and X.Wang designed the study, H.Wu, Q.Liu and X.Wang, X.Mou, C.He, and L.Tan designed the experiments, [movie, Chinese text…], X.Mou, C.He and L.Tan, H.Liang and J.Zhang conducted the experiments, X.Mou, C.He, L.Tan, H.Liang, J.Zhang and J.Yu analyzed the results. X.Mou, C.He and L.Tan wrote the first draft. All authors checked the code, wrote the manuscript, reviewed the manuscript, and approved the final manuscript.

